# Quantitative single-cell splicing analysis reveals an ‘economy of scale’ filter for gene expression

**DOI:** 10.1101/457432

**Authors:** Fangyuan Ding, Michael B. Elowitz

## Abstract

In eukaryotic cells, splicing affects the fate of each pre-mRNA transcript, helping to determine whether it is ultimately processed into an mRNA, or degraded. The efficiency of splicing plays a key role in gene expression. However, because it depends on the levels of multiple isoforms at the same transcriptional active site (TAS) in the same cell, splicing efficiency has been challenging to measure. Here, we introduce a quantitative single-molecule FISH-based method that enables determination of the absolute abundances of distinct RNA isoforms at individual TASs. Using this method, we discovered that splicing efficiency behaves in an unexpected ‘economy of scale’ manner, increasing, rather than decreasing, with gene expression levels, opposite to a standard enzymatic process. This behavior could result from an observed correlation between splicing efficiency and spatial proximity to nuclear speckles. Economy of scale splicing represents a non-linear filter that amplifies the expression of genes when they are more strongly transcribed. This method will help to reveal the roles of splicing in the quantitative control of gene expression.

## Introduction

In eukaryotes, most genes undergo co-transcriptional splicing^1–3^. During splicing, nascent pre-mRNAs are processed to remove introns and to include or exclude exons^4^, generating multiple isoforms, with distinct fates, including functional mRNAs and degradation products^5–7^. The ubiquitousness of splicing, and its tight coupling with transcription^1–3^ suggest that splicing could pay additional roles beyond diversification of protein products per se. In particular, it could act as a signal processing filter, modulating the amount of mature mRNA depending on the rate of transcription, or other parameters. In general, only a fraction of transcribed RNA is productively spliced to enable subsequent translation or other functions. Unspliced RNA is predominantly retained in the nucleus and degraded, through a quality control mechanism^8,9^. A critical feature of splicing is its efficiency, defined as the ratio of spliced RNA to total transcribed RNA (Figure 1a, SI text). In general, it has remained unclear how splicing efficiency depends on the gene expression level, and therefore what type of filter, if any, splicing provides.

**Figure 1:**
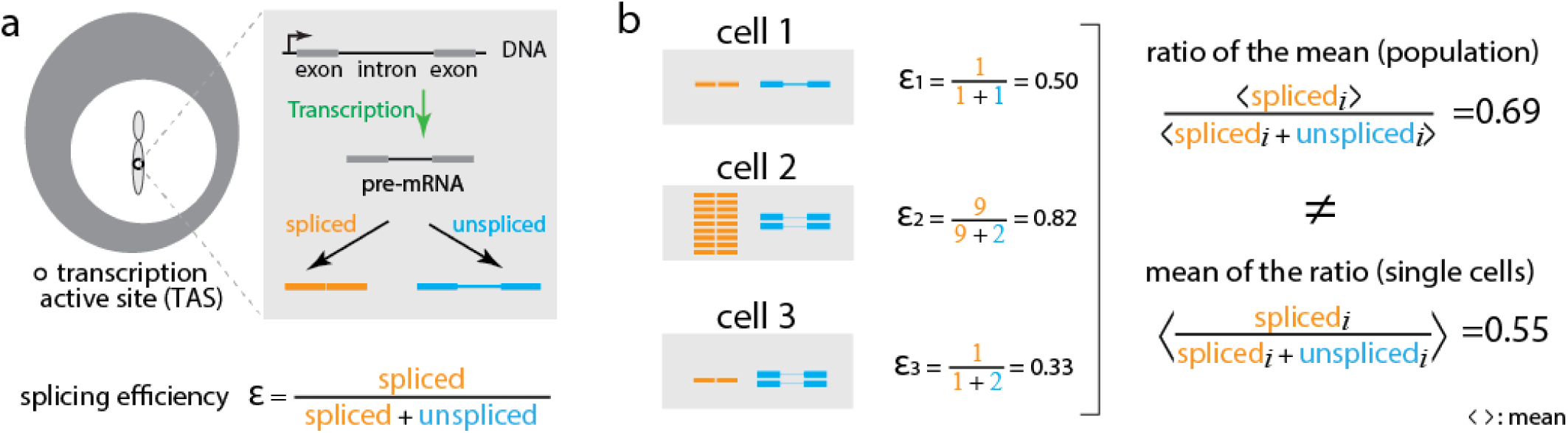
Splicing efficiency requires single cell measurement. (a) Splicing occurs co-transcriptionally at the TAS (white circle on chromosome). Each pre-mRNA molecule (gray) can be spliced to remove introns (orange) or left unspliced (blue). Splicing efficiency is the ratio between the number of spliced transcripts and the total number of transcripts at the TAS (equation, bottom). (b) Averaging over heterogeneous cells can distort splice efficiency. We consider three hypothetical cells with different numbers of spliced (orange) and unspliced (blue) transcripts. The mean of the splicing efficiency measured in each cell individually does not equal the ratio of the mean number of spliced transcripts and the mean number of total transcripts at the population level. Thus, splicing efficiency should ideally be measured in individual cells.

In general, splicing efficiency could behave in three qualitatively distinct ways. The simplest possibility is that splicing efficiency is determined by sequence features and concentrations of splicing machinery, and therefore constant for a given gene in a given cell state. However, as an enzymatic process, splicing could in principle decline in efficiency at high substrate (pre-mRNA) concentrations. This ‘diminishing returns’ behavior would tend to disproportionately suppress processing at higher expression levels relative to lower levels. The final formal possibility, which is not expected under conventional models of splicing, is that splicing efficiency could increase with transcription level, in an ‘economy of scale’ fashion, and thereby disproportionately enhance the expression of more highly expressed genes. Accurate measurements of splicing efficiency across transcription levels are needed to discriminate among these or more complex behaviors.

Several approaches have been used to measure splicing efficiency^10–14^. Genome-wide shot-gun high-throughput sequencing provides a way to compute splicing efficiency by comparing the number of reads at unspliced regions to the total number of reads in constitutive exon regions of the same gene. However, much splicing occurs co-transcriptionally^15^ and different RNA species have different lifetimes once released from the TAS. Because they do not discriminate RNAs at the TAS from RNAs at other sites, these methods can distort quantification of splice ratios. To circumvent this problem, more recent studies have performed nascent-RNAseq^16,17^ to measure co-transcriptional splicing efficiency at the TAS. These approaches have been powerful and informative. However, they necessarily average over individual cells.

Averaging over a heterogeneous cell population can, by itself, distort splicing efficiency. The key issue is that splicing efficiency is an inherently ratiometric quantity. To determine the mean splicing efficiency across a cell population, one would ideally calculate the ratio of spliced to total transcripts in each cell individually and then average this quantity over the cell population. However, because the mean of a ratio is not, in general, equal to the ratio of a mean, splicing efficiency measured from single cells does not match the population average (Figure 1b). More specifically, population-average measurements systematically underweight contributions from lower expressing cells relative to higher expression cells. For a bursty process like gene expression^18–21^, this effect can be strong.

Several previous studies have sought to analyze splicing efficiency in single cells^19,22^. Pioneering studies engineered binding sites for the MS2 and PP7 RNA-binding proteins to fluorescently label individual transcripts in live cells^23,24^. This approach enabled simultaneous analysis of splicing and transcriptional kinetics in individual cells. However, it cannot be used on endogenous (unmodified) transcripts, and insertion of binding sites could potentially perturb the splicing dynamics.

Here, we report a method for quantitative single-cell measurement of splicing efficiency based on single-molecule fluorescence in-situ hybridization (smFISH). The method measures splicing efficiency at transcriptional active sites in individual cells. In contrast to smFISH methods based on counting the number of distinct molecules, appearing as fluorescent dots in images^25–27^, here we quantify dot intensity at the TAS. For accurate quantitation, we developed methods for unbiased intensity comparisons between channels and adapted a method from astrophysics for estimating stellar luminosities in crowded star fields^28^.

Contrary to the classic enzyme-substrate Michaelis-Menten model, splicing efficiency increased, rather than decreased, with increasing levels of gene expression, in an ‘economy of scale’ fashion. Increased transcription also correlated with spatial proximity to speckles, suggesting a mechanism for economy of scale based on spatial clustering. A mathematical model based on this observation shows how economy of scale splicing could emerge if enzyme availability increases with substrate (pre-mRNA) concentration. Together, these results enable quantitative analysis of splicing in single cells and reveal a new role for splicing as a gene expression filter.

## Results

### Intron and exon probe smFISH sets identify distinct RNA species

We set out to measure splicing efficiency by quantifying the relative levels of different isoforms at the same TAS, across a range of expression levels. As a model system, we used the RG6 mini-gene, whose splicing behavior was previously characterized using fluorescent proteins^29^. To enable regulation of transcription, we site-specifically integrated the mini-gene under the control of a dox-inducible promoter into HEK293 cells. To measure the splicing efficiency at the TAS, we designed three smFISH probe sets. The intron probe set targeted the spliceable constitutive intron, and thus measured the number of unspliced transcripts. The other two probe sets, denoted Exon1 and Exon2, targeted constitutive exons, measuring the number of total transcripts (Figure 2a). The use of two redundant exon probe sets facilitates subsequent analysis (see below). We cultured cells under standard conditions and then fixed and imaged the cells using all three smFISH probe sets (SI Methods and Materials).

**Figure 2:**
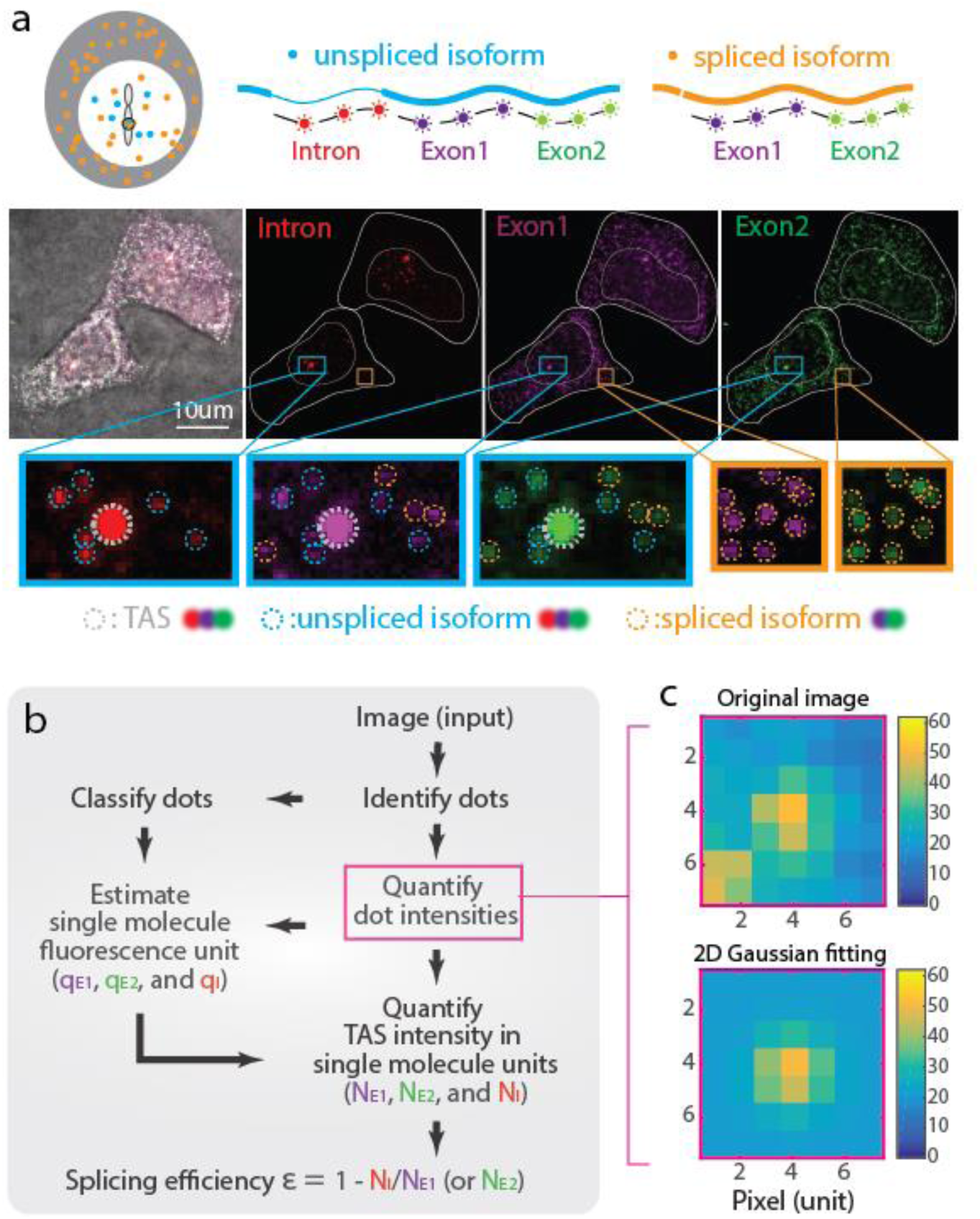
Experimental method for quantifying splicing efficiency at the TAS in individual cells. (a) We designed three smFISH probe sets targeting one spliceable intron (red), and two constitutive exons (purple, green). In this image of two cells, staining by each probe set is shown in the corresponding color channel, superimposed on a brightfield image (grayscale). Identified smFISH dots are circled in the zoomed insets, with TAS in gray, unspliced isoform (dots co-localized in Intron, Exon1, and Exon2 channels) in blue, and spliced isoform (dots co-localized in Exon1 and Exon2 channels) in orange. (b) Workflow for quantifying N_E1_, N_E2_ and N_I_, the number of transcripts observed within the TAS in the exon1, exon2, and intron channels using corresponding single molecule fluorescence units, q,_E1_, q_E2_, and q_I_, respectively. (See also Figure S1a.) (c) Iterative fitting of individual dot intensities. Using a method from stellar photometry of crowded star fields, we iteratively remove fluorescence from adjacent dots to produce a single dot image (bottom) from an initial image with multiple dots (top) (see SI Text and Figure S1c for more detail).

We observed several types of smFISH dots that could be classified by their intensity and probe binding patterns. The first type, representing the TAS, consisted of one or two bright dots per cell that appeared in all three channels in the nucleus (Figure 2a and SI Text). The second type consisted of many scattered dots of lower intensity that appeared in both the nucleus and cytoplasm corresponding to single transcripts (Figure 2a). Using co-localization of different probe sets enabled further classification of dots. All RNAs were labeled by both Exon1 and Exon2 probe sets, both in the cytoplasm and the nucleus. In contrast, unspliced isoforms were labeled in all three channels, but appeared only in the nucleus (Figure 2a). Although we targeted a constitutively spliced intron, we also observed released unspliced molecules, consistent with imperfect splicing efficiencies (Figure 2a, ‘unspliced isoform’). These transcripts could result from a failure to complete splicing prior to transcriptional termination, or from competition among transcripts for limited levels of splicing machinery^30^. Together, these probe sets allowed identification of multiple distinct molecular species both at and away from the TAS.

### Determining single transcript intensity units

Because individual transcript molecules cannot be spatially resolved at the TAS, we developed an intensity-based transcript-counting procedure to quantify the number of transcripts of each species from fluorescence intensities of each probe set (Figure 2b, Figure S1a and SI Text). Briefly, we first used the Poisson-distributed individual transcript intensities to obtain the single molecule fluorescence intensity calibration units in each color channel, denoted qE1, qE2, and qI for the two exon and one intron channels, respectively. Then, we used this calibration to quantify the number of copies of exonic (N_E1_, N_E2_) and intronic (NI) targets at the TAS, in molecular units.

In this procedure, the values of q_n_ (n = E1, E2, or I) are estimated from the distribution of intensities of single-molecule non-TAS smFISH dots in each channel. However, dot identification can be inconsistent between channels due to differences in signal-to-background levels, differences in binding efficiencies between probe sets, and differences in the fluorescence properties of each fluorophore, among other issues. These factors produce systematic differences in sensitivity between channels that distort qn quantification.

To address this issue, we developed an unbiased dot identification procedure. We first identified candidate dots using a low (permissive) threshold that captures all foreground smFISH dots as well as some background signal (false positive dots) in each channel, and quantified the integrated intensity of each dot. To correct for fluorescence ‘contamination’ from neighboring dots, we adapted an algorithm from stellar photometry of crowded star fields^28^, which works by iteratively removing fluorescence from neighboring objects (Figure 2c and Figure S1c, SI Text).

We also performed this analysis on negative-control images lacking true smFISH dots to obtain the distribution of background dot intensities. Subtracting the background histogram from the total (foreground + background) histogram generated a corrected foreground dot intensity distribution (Figure 3a, and SI Text). TAS intensity measurements do not require accurate counting of scattered smFISH dots, as in conventional FISH^25^. This approach thus effectively sacrifices precision in individual dot identification to obtain a less biased distribution of smFISH dot intensity in each fluorescent channel. Finally, to obtain the single transcript intensity unit qn, we fit the resulting distributions with a continuous analog of the Poisson distribution (Figure 3b, and SI Text).

**Figure 3:**
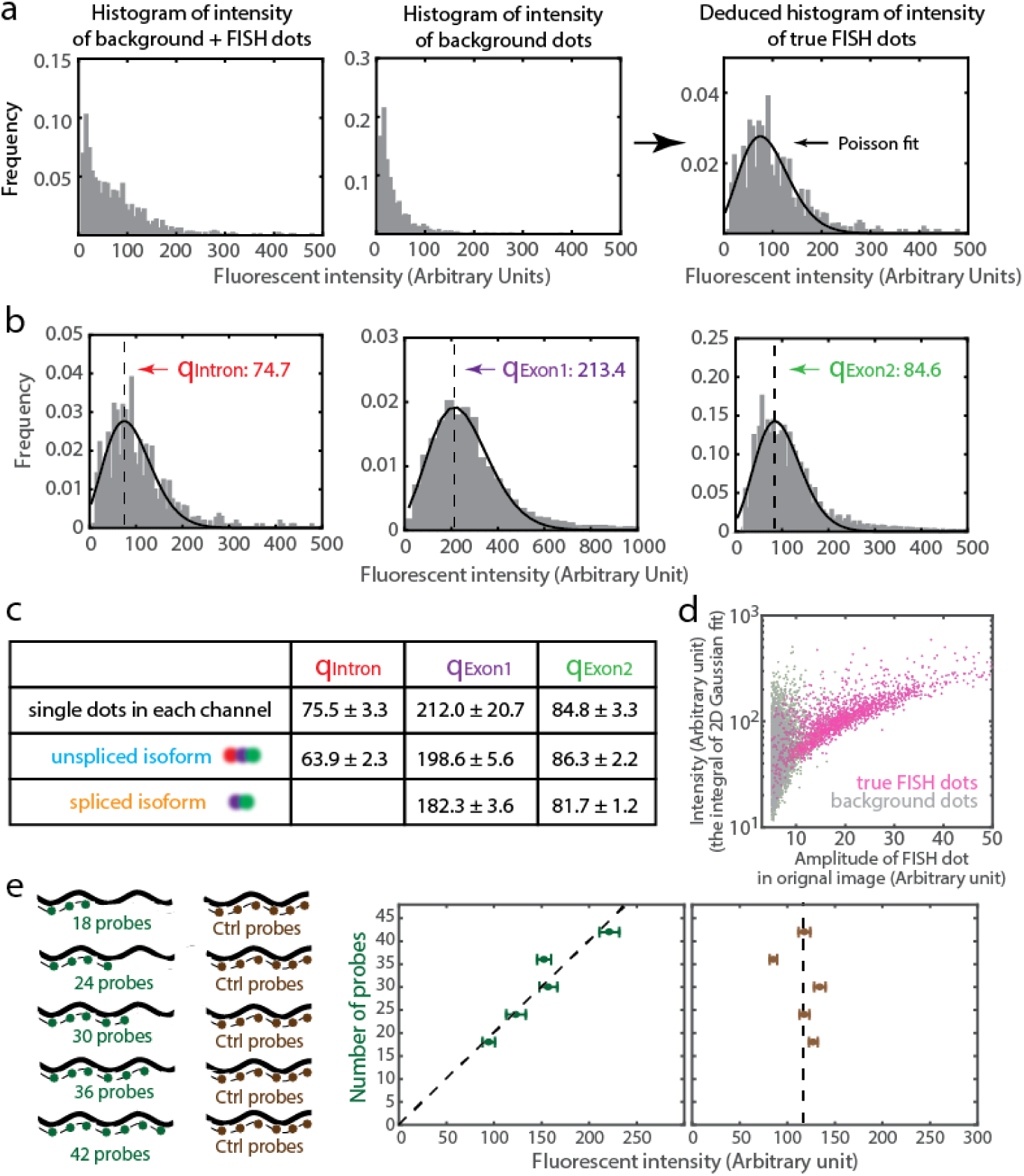
Quantification of single transcript intensity units in multiple channels. (a) The single dot transcript intensity distribution (right) can be obtained by subtracting a background distribution obtained without FISH probes (middle) from the foreground+background image obtained with FISH probes (left). (b) Applying this procedure to each channel, and fitting the resulting distributions to a Poisson distribution provides the three single transcript intensity units (indicated). (c) The measured single transcript intensity units in multiple channels are verified via dot-colocalization (Figure S1d). Different methods generate similar single molecule fluorescence units. Color is labeled as in Figure 2a. (d) The properties of background dots (gray) and identified true FISH dots (pink). Note overlap of these properties. See more dot fitting properties in Figure S2-S4. (e) The measured fluorescent intensity for staining of SDHA gene is proportional to the number of smFISH probes included (indicated numbers). A negative control (brown) uses a fixed number of probes, and displays a relatively fixed fluorescence intensity. This indicates that intensity scales linearly with number of probes.

As an additional consistency check, we used an independent method to identify foreground dots by their co-localization across multiple fluorescence channels. This approach produced similar values for q_n_ (Figure 3c, Figure S1d, SI Text). It also revealed that foreground and background dots exhibit overlapping distributions of key properties such as intensity, peak height, and peak width (Figure 3d, and Figure S2, S3, S4, SI Text), further supporting the need for histogram subtraction, above. Taken together, these results provide a simple and general method for accurately estimating the fluorescence intensity at the TAS using precisely calibrated intensity units for single transcripts.

### Fluorescence intensity increases linearly with number of probes

Because intensity quantification is critical for estimating the number of molecules at the TAS, we next asked how TAS fluorescence intensity depends on the abundance of bound probes. We designed 42 smFISH probes targeting the housekeeping gene SDHA, and mixed them into groups of 18, 24, 30, 36, and 42 probes. We then measured the fluorescence intensity of stained HEK293 cells with each group. Fluorescence dot intensities increased linearly with the number of included probes (Figure 3e). By contrast, a second set of 27 probes targeting HES1, which were labeled in a different fluorescent channel and included in all experiments as a fixed control, were constant across each condition, as expected. These results indicate that fluorescence intensity provides a linear readout of probe density at the TAS. This linearity enables one to quantify the number of transcripts (N_E1_, N_E2_, and N_I_) at the TAS in molecular units by dividing TAS fluorescence intensity by the single transcript fluorescence units (q_E1_, q_E2_, and q_I_, respectively).

### Splicing efficiency exhibits an ‘economy of scale’ behavior

Having established the method, we next used it to determine how splicing efficiency changes with transcription level in individual cells. Quantifying splicing efficiency requires comparing the number of spliced transcripts to the total number of transcripts (spliced + unspliced) at the TAS. Here, the total transcript number was represented by N_E1_ or N_E_, and the number of spliced transcripts was obtained by subtracting the number of pre-spliced transcripts NI, from NE1 or N_E2_. With these quantities, the splicing efficiency can be computed as ε_i_ = (N_Ei_ - N_I_)/N_Ei_ = 1 – N_I_/N_Ei_, where i=1 or 2 denotes either of the two exon probe sets, and εi should, in the absence of noise, be independent of i.

To cover a broad range of expression levels, we induced the Sonic Hedgehog (SHH) target gene Gli1 in 3T3 cells with varying concentrations of recombinant SHH, and analyzed cells after 48h of induction^31^. In parallel, we analyzed the synthetic spliceable RG6 mini-gene described above, induced for 3h with a range of doxycycline concentrations. After induction, we fixed cells and performed smFISH-hybridization and imaging (SI Methods and Materials).

The Gli1 and CMV promoters expressed at up to ∽20 transcripts per TAS. Altogether, we analyzed ∽3000 (Gli1, Figure 4a) and ∽1000 (RG6, Figure S5a) active sites in single cells, and computed the geometric mean for each condition (SI text), as well as the splicing efficiency in each cell (Figure 4b, and Figure S5b). For both genes, transcription level and splicing efficiency were heterogeneous, even at a single induction level^18,32^. This variability likely reflects both transcriptional bursting and other sources of biological variation^32^, as well as measurement errors from stochastic binding of probes and other sources (see SI text for more details).

**Figure 4:**
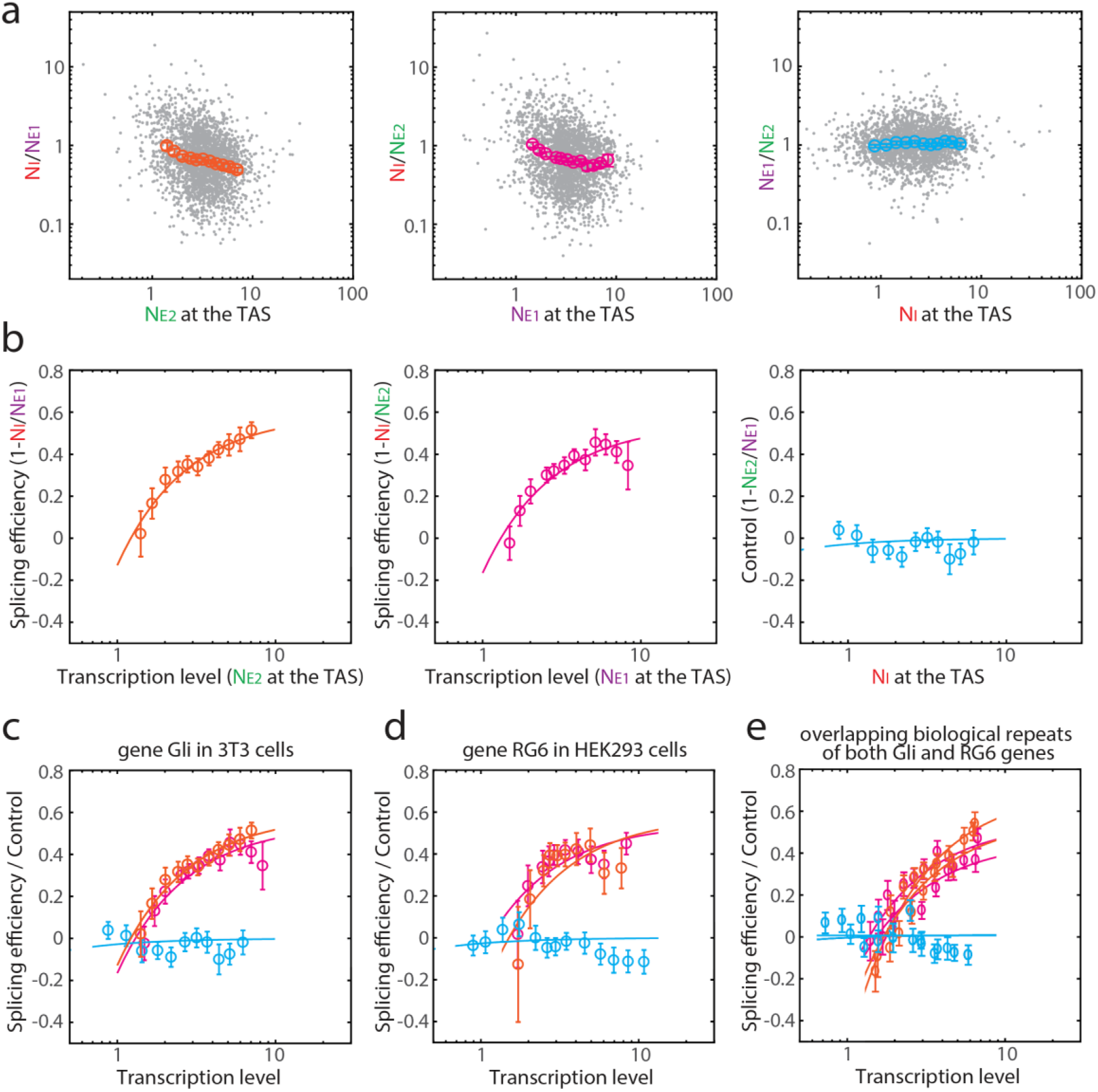
Splicing efficiency increases with transcript level, exhibiting ‘economy of scale’ behavior, for the two tested genes, Gli1 induced by Shh in 3T3 cells and the Tet-off mini-gene RG6 induced by dox in HEK293 cells. (a) Raw data of the number of transcripts measured by our methods. Each dot (gray) in the plot is one measurement from a single TAS. The number of TAS measured is ∽3000 for Gli1 (and ∽1000 for RG6 in Figure S5). Geometric means are shown in orange/pink/blue. Error bars represent standard error of the mean. (b) Based on (a), we find splicing efficiency (1-N_I_/N_E1_, or 1-N_I_/N_E2_) increases with transcription level (orange and pink), while control (1-N_E1_/N_E2_, or 1-N_E2_/N_E1_) measurements remained constant (blue). Solid lines are guides to the eye to highlight the ‘economy of scale’ behavior. The ‘economy of scale’ mathematical model is discussed in the main text and the corresponding Michaelis-Menten model fitting on the raw data were shown in Figure S9c. (c) Overlapped curves from (b). (d) Overlapped curves of the synthetic gene RG6. (e) Overlapped curves from biological repeats for Gli1 and RG6. All of them showed the repeatability of ‘economy of scale’ observation.

While individual cells were variable, the mean splicing efficiency systematically increased with gene expression level. This ‘economy of scale’ behavior occurred for both genes. It was robust to experimental conditions, such as the strength and duration of induction. It was also robust to data analysis parameters (see SI text). Additionally, splicing efficiency increased with transcription level in a similar pattern for both genes (Figure 4c, 4d, 4e), reaching 80% of its maximum value at ∼3.5 transcripts per TAS.

We ruled out potential artifacts that could appear to generate this ‘economy of scale’ behavior. For example, because imaging occurs at a fixed point in time, images can in general capture incomplete transcription events. If incomplete transcription were expression level-dependent, it would alter the two exon probe ratio, N_E1_/N_E2_, from its ideal value of 1. However, this ratio showed no systematic dependence on transcription level (Figure 4 and Figure S5). Additionally, to rule out potential misclassification of individual transcripts as the TAS, we used single-molecule DNA FISH to independently identify the TAS (Figure S6). Finally, analysis of two independent transcription level measurements (N_E1_ and N_E2_) enabled us to compute splicing efficiency and transcription rate using distinct exon readouts, avoiding a potentially spurious correlation between transcription level and splicing efficiency due to the appearance of NE1 in the expression for splicing efficiency, (1-N_I_/N_E1_). (Note that without the second exon probe, hardness-ratios correction methods^33,34^ from Astrophysics could also help address this issue (Figure S7 and SI text)). Together, these results support the validity of the economy of scale observation.

### A mathematical model of ‘economy of scale’ splicing

How can an enzymatic process such as splicing produce ‘economy of scale’ behavior? In the classic Michaelis-Menten model (Figure 5a and SI text) reaction efficiency declines monotonically with increasing substrate concentration (Figure 5b, black curve), producing the opposite ‘diminishing returns’ behavior.

**Figure 5:**
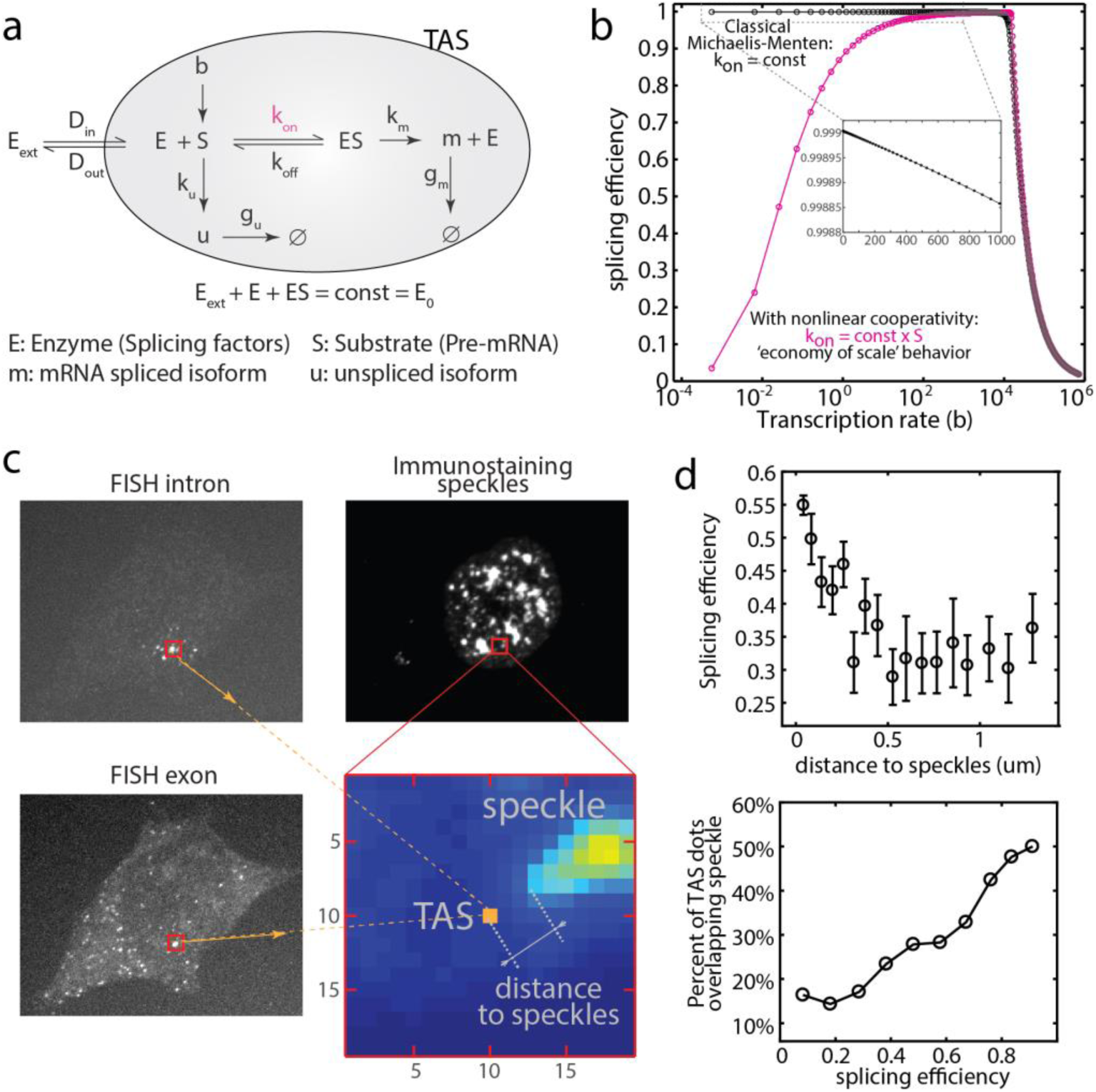
Potential model for ‘economy of scale’ behavior. (a) In a simple scheme based on the classical Michaelis-Menten model, splicing factors, collectively denoted E, bind to pre-mRNA, denoted S, at a rate k_on_ to form an enzyme-substrate complex, ES, which can unbind, at rate k_off_, or transform into a mature mRNA, m, at rate k_m_, releasing the enzyme. k_u_ denotes the production rate of unspliced RNA, denoted u, g_u_ and g_m_ denote the rate at which unspliced RNA or mRNA, respectively, are degraded or released from the TAS (shaded gray area). E_0_ denotes the total concentration of enzyme. (b) In the classical Michaelis-Menten model, splicing efficiency declines monotonically with transcription rate, in a “diminishing returns” manner (black). A variant of the model in which k_on_ is proportional to S, rather than constant, generates ‘economy of scale’ behavior (pink). Note that at higher transcription rates, splicing efficiency declines due to enzyme saturation. These ‘diminishing returns’ and ‘economy of scale’ behaviors were parameter independent (Figure S9b and SI text). (c) We measured the distance between TAS and its nearest speckle via smFISH and immunostaining in the same cell. FISH probes were designed as in Figure 2a. We used SC35 antibody to immunostain the speckles. The zoomed-in area was marked in red. smFISH defined the TAS location (orange), and immunostaining intensity showed the presence of speckles (Figure S8). (d) Splicing efficiency increases with proximity to speckles (top figure). TAS with higher splicing efficiency has greater probability of overlap with speckles (bottom figure). See raw data in Figure S8b.

One limitation of the simple Michaelis-Menten model is that it does not account for the inhomogeneous concentration of splicing machinery in dynamic interchromatin granule clusters called nuclear speckles^35–37^. Previous work has shown that more highly transcribed genes are closer to speckles^36,38^. To analyze the relationship between spatial organization and splicing efficiency, we analyzed splicing efficiency of RG6 as described above, and also performed immunostaining to detect the splicing factor SC35 in the same cell (Figure 5c, SI Methods and Materials). We then quantified the distance from each TAS to its nearest speckle (Figure S8a and SI Text), and plotted the results as a function of splicing efficiency. This analysis revealed a correlation between splicing efficiency, transcription level, and the proximity of the TAS to the splicing speckle (Figure 5d and Figure S8b).

Together, these results suggest the hypothesis that stronger expression could increase the proximity of a gene to a speckle, which in turn could increase the availability of splicing machinery. To incorporate these effects into a modified version of the model, we allowed the rate of pre-mRNA binding to splicing machinery, k_on_, to increase with the concentration of pre-mRNA (Figures 5. This simple modification generated economy of scale behavior at lower expression levels (Figure 5b, pink curve, and schemed in Figure S9a), switching to diminishing returns at higher expression levels as the splicing machinery eventually saturates (see additional mechanisms in the SI Text). These results show that a positive correlation between expression level and speckle accessibility could qualitatively explain ‘economy of scale’ splicing behavior.

## Discussion

Here, we introduced a quantitative imaging-based method to measure splicing efficiency in single cells, and used it to characterize the dependence of splicing efficiency on transcription level. It enables accurate intensity quantification of smFISH data to allow direct comparison of intensities of multiple channels at the same site. Although we focused on quantifying the splicing efficiency of constitutively spliced introns, the pipeline presented here can be extended to alternative splicing by incorporation of additional fluorescence channels.

The observed ‘economy of scale’ behavior is opposite to the ‘diminishing returns’ behavior one would expect from a standard enzymatic process. Mechanistically, it could reflect a disproportionate allocation of the shared splicing machinery ‘resource’ to more highly expressed genes. In fact, previous work has shown that different genes can effectively compete for splicing machinery inside the nucleus^30^, with more weakly expressed genes receiving less access to splicing factors^36,38.^ This allocation of splicing resources could optimize the total amount of splicing that can be achieved by a fixed abundance of splicing components^39^.

Functionally, economy of scale acts as a non-linear filter within the overall gene regulation process, enhancing more strongly expressed genes (Figure 6a). In principle, this filter could impact many cellular regulation processes. For instance, it could help prevent pervasive low-level transcription^40^ from inappropriately propagating to protein synthesis. It could also amplify, suppress, or reshape the mRNA distribution, depending on the underlying distribution of transcription levels (Figure 6b and 6c). Thus, it could play an active role in controlling variation across a population of cells^41,42.^

**Figure 6:**
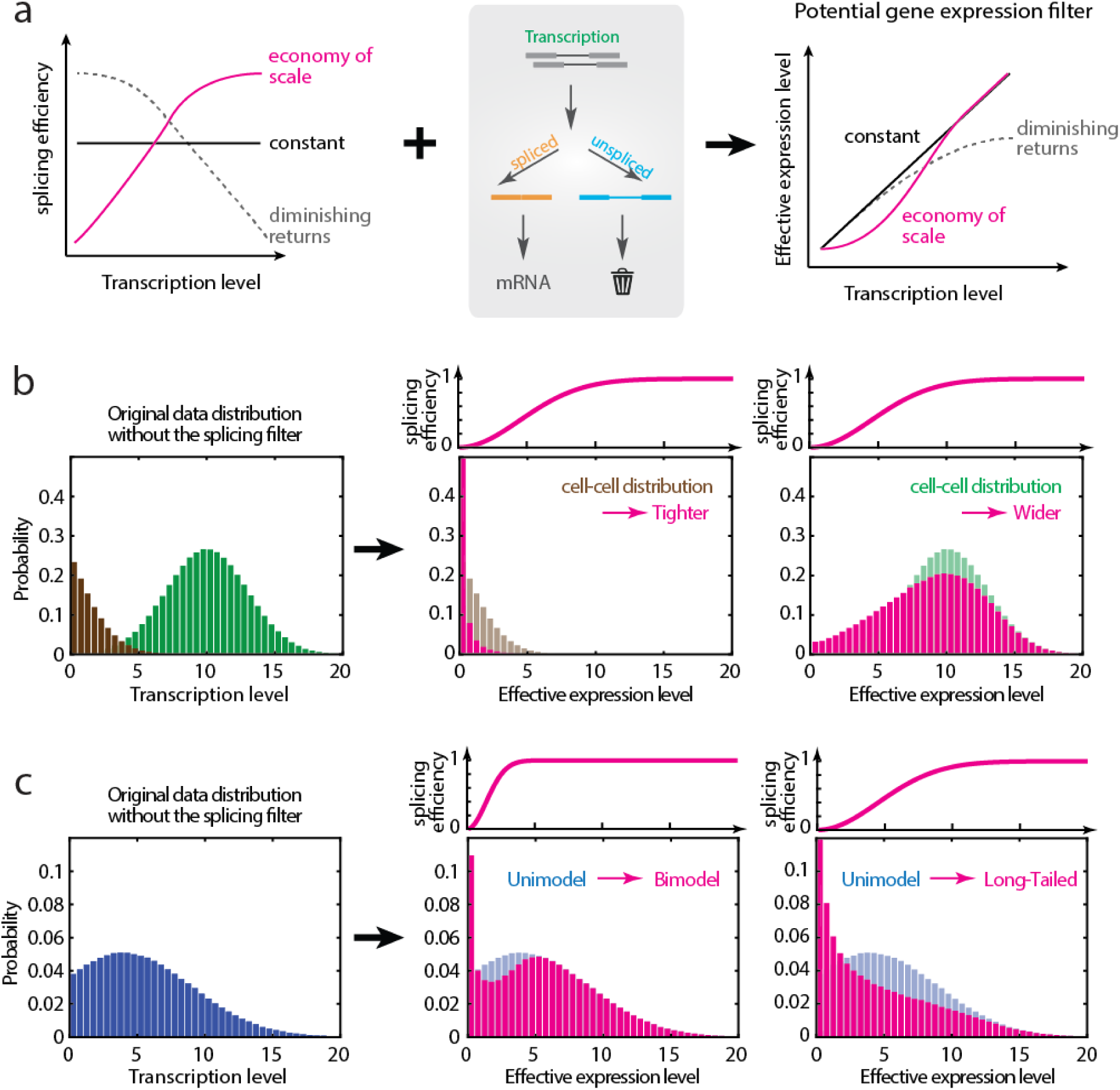
Splicing can act as a gene expression ‘filter.’ (a) Different possible relationships between splicing efficiency and transcription level can generate different type of gene expression filter. ‘Economy of scale’ (pink) and ‘diminishing returns’ (gray) are high-pass and low-pass filters, respectively. (b) The ‘economy of scale’ filter can narrow or broaden the distribution of mRNA levels depending on the underlying distribution of mRNA expression levels. The filter (pink) preserves higher transcription levels (green) but suppresses lower transcription levels (brown). (c) Depending on its parameters, the ‘economy of scale’ filter can change the shape of an expression distribution from unimodal (blue) to bimodal (pink, center plot) or long-tailed (pink, right hand plot).

There are several limitations of the present method. One is that it focuses on co-transcriptional splicing. Potential regulation through post-transcriptional splicing processes^24,43^ is not captured. Second, because the method relies on direct binding of probes, occlusion of probe binding sites by bound proteins or secondary structure of the target RNA could in principle affect quantitation. Third, the post-splicing residence times for different products at the TAS are unknown. Their relative values affect the absolute magnitude of the measured efficiency, but not its dependence on transcription level. In principle, however, the dependence of splicing efficiency on transcription rate could be distorted if the residence times for different molecules depend in different ways on transcription rate (SI text).

Because it operates quantitatively in single cells with subcellular resolution, this method should provide insight into kinetic features of the splicing mechanism. For example, by simultaneously imaging the spatial locations of splicing regulatory factors such as lncRNA alongside their target genes, it could enable one to determine how these factors affect splicing efficiency^44–48^. Using additional fluorescent channels, it could also allow analysis of correlations in splicing between neighboring genes, and enable comparison splicing efficiency between alleles of a gene within a single cell.

Further development of the method could address additional questions. For example, recent work has shown how comparison of nascent transcript levels with total transcript levels can provide information on dynamic changes in expression (RNA ‘velocities’) from single time-point snapshots^49^. These approaches could be combined with the analysis shown here to provide dynamic information on the relation of splicing to transcriptional bursting. Finally, by combining these approaches with sequential hybridization and barcoding techniques^44,45,50,^ this method could enable genome-wide analysis of splicing efficiency in a single cell.

## Acknowledgements

The work was funded by a Fellowship from the Schlumberger Foundation, by the Gordon and Betty Moore Foundation through Grant GBMF2809 to the Caltech Programmable Molecular Technology Initiative and the Institute for Collaborative Biotechnologies through grant W911NF-09-0001 from the U.S. Army Research Office. The content of the information does not necessarily reflect the position or the policy of the Government, and no official endorsement should be inferred. M.B.E. is a Howard Hughes Medical Institute Investigator.

